# Resistance screening of uncharacterized *Lupinus angustifolius* accessions towards *Fusarium oxysporum f. sp. lupini*

**DOI:** 10.1101/2025.04.30.651460

**Authors:** Camila Rayo, Elias Messner, Karina Tokina, Thomas Svoboda

## Abstract

Lupins, which belong to the family of fabaceae, are plants with a high nutritional potential due to their high protein content, symbiotic nitrogen fixation and environmental adaptability. Yet, plant pathogens including *Fusairum oxysporum f. sp. lupini* are a considerable problem in lupin cultivation as they can cause considerable yield reduction. To find *Lupinus angustifolius* accessions with enhanced resistance towards Fusarium infection, we screened 20 yet uncharacterized accessions in a root infection assay. The infected roots were harvested four days post inoculation, followed by total DNA extraction. The rate of infection was estimated by calculating the relative amount of lupin and fungal DNA based on the respective qPCR cq-values. After screening of all 20 accessions, we identified L26 as the most resistant and L49 as the most susceptible accession. To verify this, infection assays of a longer time period of 15 days were performed with these two accessions. The initial results were confirmed as more severe symptoms were observed on L49. To better characterize the infection dynamics, a time series of the infection process was set up and infected roots were evaluated after one, two three and four days. These results showed that already after two days post inoculation there is a significant difference regarding infection levels between L26 and L49. Further characterizations will provide valuable information on the genes responsible for the different resistance levels.

## Introduction

*Lupinus ssp*. is a fast-growing annual grain from the fabaceae family among which *Lupinus angustifolius*, also known as narrow-leafed lupin (NLL), *Lupinus albus, Lupinus mutabilis* and *Lupinus luteus* are commercially cultivated. Lupins have a high nutritional profile in terms of lipids (up to 18% dry weight) and proteins (up to 44% dry weight). Additionally, lupins produce different phytochemicals like flavonoids, alkaloids, triterpenoids and phenolic acids which have been used for medical purposes (Ishaq et al. 2022). Quinolizidine alkaloids have different affinities to various acetylcholine receptor subunits resulting in species-specific toxicities and clinical diseases (Panter, James, and Gardner 1999). Hence in food and feed production quinolizidine alkaloids are not desired due to their bitter taste and detrimental effects. This led to breeding of “sweet” lupin accessions which do not have a sweet taste but produce significantly lower amounts of bitter quinolizidine alkaloids. It has been demonstrated in frame of the domestication of the white lupin that a single nucleotide mutation in the pauper locus is sufficient to block the production of toxic alkaloids due to strong impairment of acetyltransferase activity (Mancinotti et al. 2023).

A relevant trade-off for the low alkaloid content, however, is a higher susceptibility of these lines towards different fungal pathogens which has been correlated to a function of these alkaloids as antifungal compounds (Zamora-Natera et al. 2008; Cely-Veloza et al. 2023). It has been demonstrated in seed infection assays of *Lupinus luteus* with *Fusarium oxysporum f. sp. lupini* that the initial defense response of the plant is mediated through the production of isoflavonoids (Morkunas et al. 2010). On the other side, quinolizidine alkaloid biosynthesis does not seem to have an impact on anthracnose resistance thus making other alkaloids equally important for pathogen resistance (Czepiel et al. 2021). While for anthracnose resistance the LanrBo locus has been identified in NLL (Fischer et al. 2015), there has no gene or gene cluster been described yet for Fusarium resistance. *Fusarium oxysporum f. sp lupini* (Fol) is a plant pathogen responsible for causing Fusarium wilt. Fol produces durable chlamydospores which can overwinter in the soil and can be spread by wind and water. The early symptoms of fusarium wilt include yellowing of lower leaves and stunting of the plant. It further causes vascular discoloration and the entire plant may die (Dean et al. 2012). Beside the yield reduction, *Fusarium oxysporum* produces several toxic secondary metabolites which are harmful for humans and animals, including fusarins, fusaric acid and moniliformin (Perincherry, Lalak-Kańczugowska, and Stępień 2019).

For the above-mentioned reasons, cultivation of plants with a higher resistance against Fol is desired. In this context we are screening yet uncharacterized NLL accessions in terms of Fol resistance which are originating from the Vavilov Institute of General Genetics and maintained by a commercial lupin breeder. Here we report the establishment of a Fol screening method with lupin plants including optimal conditions for Fusarium infection and a DNA-based determination of resistance levels. With this method we analyzed a first set of 20 yet uncharacterized accessions and found one to be highly resistant against Fol infection in several independent assays.

## Material and Methods

### Seed sterilization and germination

For sterilization of the lupin seeds, the seeds were rinsed with 75% ethanol for one minute followed by immersion in a 10% NaClO solution for 15 minutes, shaking gently. After discarding NaClO, the seeds were washed four times with water. For germination, a sterile filter paper was put into a petri dish (90mm x 15 mm), soaked with 8 ml sterile water and ten seeds were put on one plate. The plates were sealed with parafilm and incubated at 20°C with a day/night cycle of 16h/8h at 60% humidity.

### Fusarium cultivation

In this study *Fusarium oxysporum* f. sp. *lupini* Snyder et Hansen (ATCC 18776) strain was obtained from the American Type Culture Collection (ATCC) was used. Spores were produced on V8 media (for 800 ml: 160 ml V8 juice (Campbell Soup Co.), 2.4 g CaCO_3_, 16 g Agar-agar). Therefore, a freezer culture was evenly distributed on the plate and incubated at 20°C with 12h/12h light/dark.

### Seedling infection

For seedling infection, the spores were harvested from the plate, counted and diluted to a final concentration of 5*10^4^ spores/ml. The roots of the lupins germinated for three days were put in the spore solution and incubated for 30 minutes. Subsequently, the infected and mock treated plants were put in a 25×200 mm DURAN test tube containing 20 ml of a 1% agar followed by incubation at 20°C, 60% humidity, with a 16h/8h day/night cycle for four days. After four days the roots were harvested by cutting them below the cotyledons followed by freezing in liquid nitrogen and lyophilization.

### qPCR

DNA was extracted according to (Cenis 1992). For qPCR a 1:5 dilution of the extracted DNAs was prepared. In this study already established primer pairs were used, PDF2 for *Lupinus angustifolius* and Foxy for *Fusarium oxysporum* (Taylor et al. 2016; Rocha et al. 2023). The qPCRs were prepared according to the Luna® Universal qPCR Master Mix instructions in a total volume of 10 µl. The cycle started with 1 minute at 95°C initial denaturation followed by 40 cycles of 95°C, 15 seconds and 55°C for 30 seconds followed by fluorescence detection. After the cycles were finished a melting curve was generated by gradually increasing the temperature from 60°C to 95°C and fluorescence detection every 0.5°C.

## Results

### Method establishment

For the establishment of the method several parameters had to be evaluated and optimized. We started with testing primer pairs which were used to quantify the amount of lupin and Fusarium DNA, respectively, present in the total DNA isolated from the samples. For that purpose, genomic DNA was extracted from three random NLL accessions as well as from Fol independently. PCRs were set up to check specificity, i.e. if only the target region is amplified with the respective primer pair exclusively in the designated organism. The control gel showed that amplification of the respective region was successful and no unspecific PCR products occurred (Figure 1A). In the next step, lupin roots were infected and the roots were harvested after four days. Total DNA was extracted and qPCR was performed with the aim to check the specificity of the primers in mixed samples. The melting curves showed clearly that only one fragment was amplified (Figure 1B).

**Figure 1:**
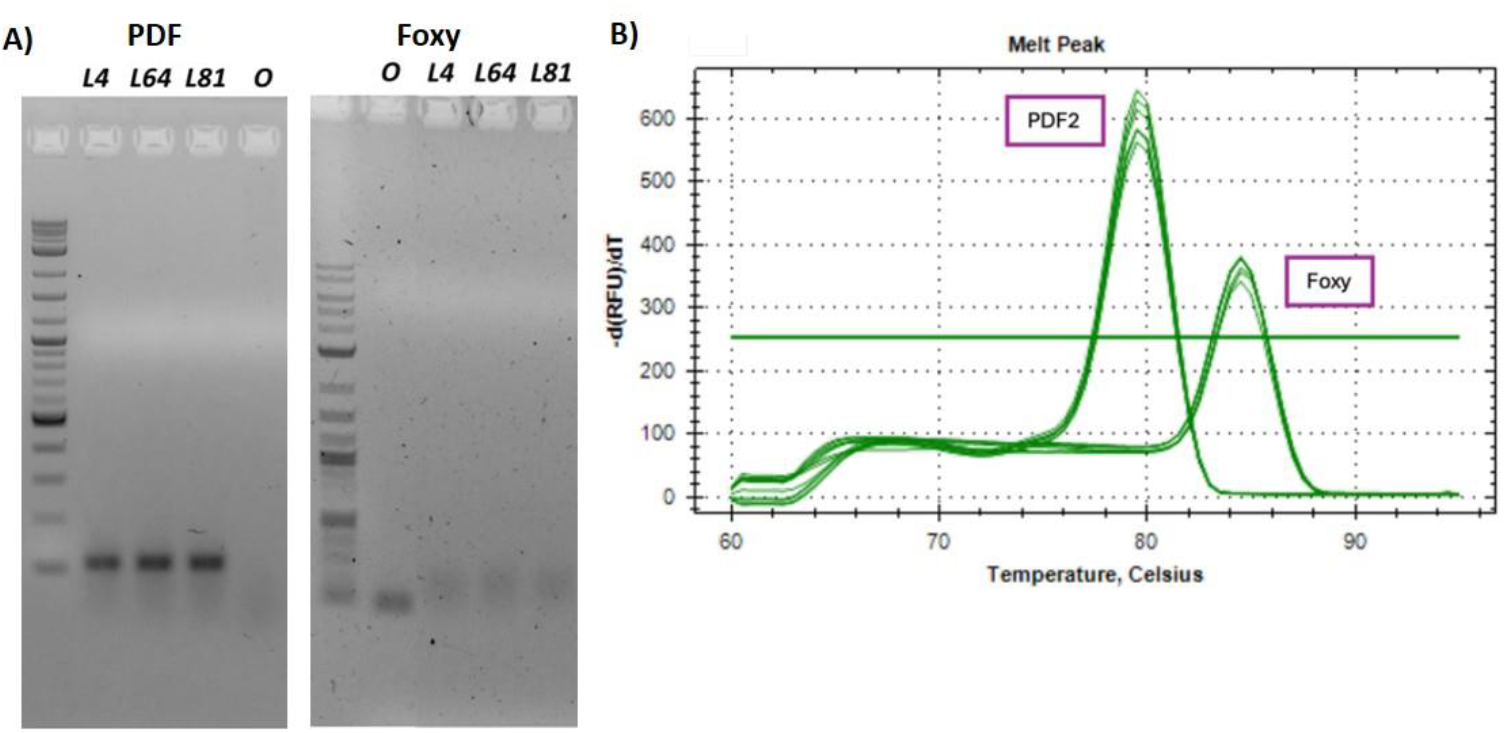
Validation of the PCR conditions A) PCR with pure DNAs to test the specificity of the primers (PDF = 109 bp (left), Foxy = 98 bp (right)); O = Fol DNA used; L4, L64, L81 = DNA of the respective lupin accession used B) melting curves of PDF2 and Foxy primers after qPCR of infected lupins; the lowest band of the ladder indicates 100 bp

This confirmed that the primers were highly specific with pure DNA as well as in the mixed samples, and we therefore proceeded with the resistance screening of the accessions.

### Assessment of resistance levels of 20 NLL accessions towards Fol

The resistance levels of the accessions were assessed by using the ratio of the cq values, which indicate a cycle threshold where the DNA can be detected, of the lupin and the fungal primers. According to the qPCR principle, the more fungal DNA is present in the plant the lower the cq value will be. Hence a higher ratio indicates that more fungal DNA was present and thus the accession is more susceptible. In contrast, a lower ratio indicates a lower amount of fungal DNA (higher cq value) in the sample and hence a higher resistance of the tested line. Based on these values we aimed to classify the different accessions according to their resistance levels. To the determine the resistance of the different NLL accessions, the infection was performed in two biologically independent experiments with three and six independent replicates, respectively, for every accession. After harvesting and lyophilization, qPCR was performed. For the evaluation of the qPCR, the ratio of the cq values of the lupin and Fusarium amplicon were calculated.

The results of the two independent approaches were statistically evaluated. This was further evaluated using Tucky’s HSD post hoc test resulting in classification of resistance/ susceptibility levels. Among the screened accessions we identified L26 as the most resistant one and L49 as the most susceptible accession (Figure 2).

**Figure 2:**
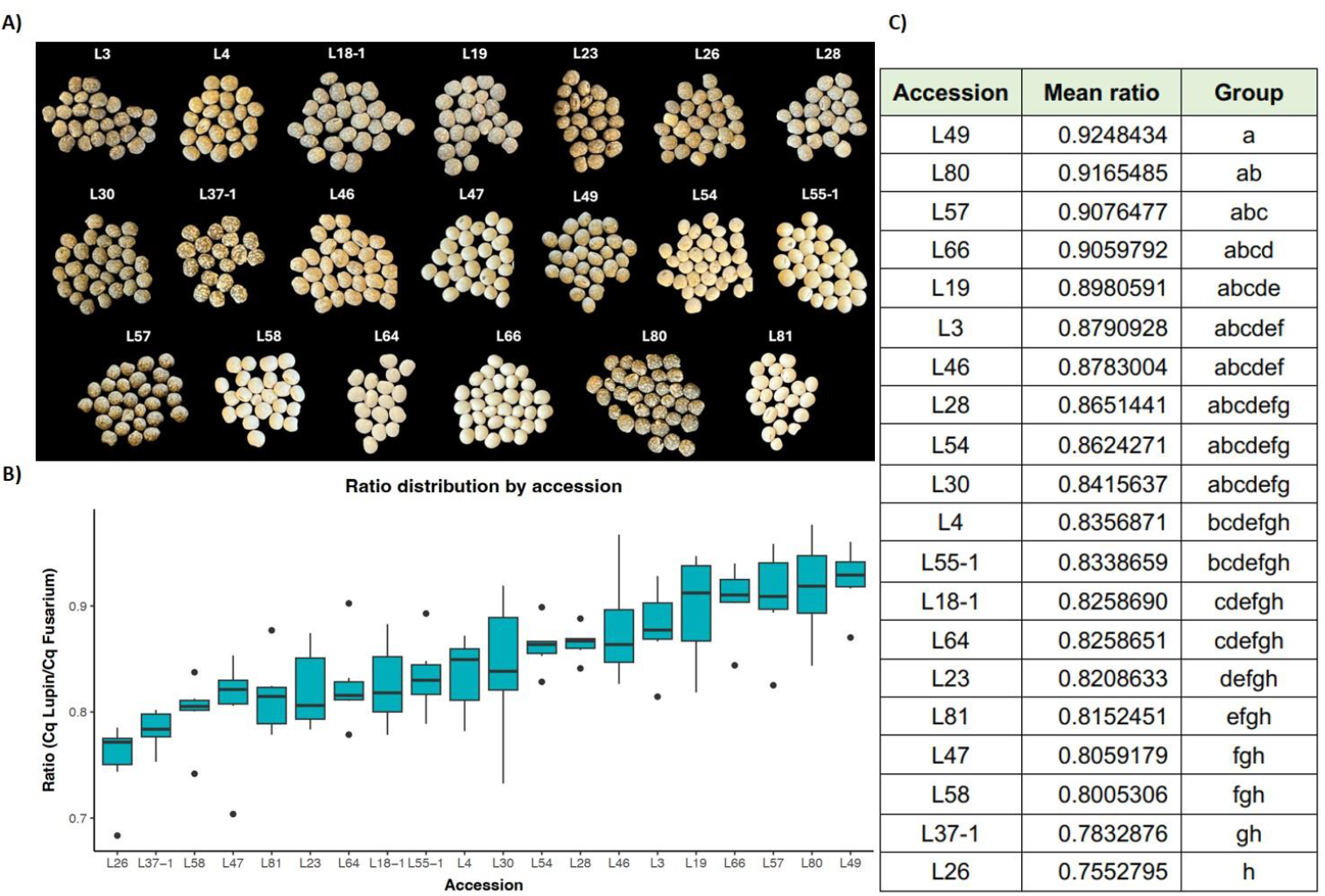
A) appearance of the seeds of the different accessions; B) boxplot of the screened NLL accessions sorted according to their resistance levels; C) mean ratios and assignment of all accessions to the respective group according to the resistance level

In the previous experimental setup, the roots were harvested after four days where phenotypically no differences were visible. To determine whether the differences in the resistance levels are phenotypically visible, an infection experiment over a longer time period of 15 days was done comparing the most susceptible (L49) and the most resistant (L26) accessions. After 15 days the phenotypic appearance of the roots and the shoot was evaluated to determine whether differences in the resistance levels determined by qPCR after 4 days also have a phenotypic impact after 15 days post infection. The symptoms observed on L26 were much less severe compared to L49 (Figure 3). While L26 only showed a slight discoloration of the root, the root of L49 was completely brown and also some mycelium was growing on the outside. Further, there is a strong difference visible regarding growth between the two accessions. While L26 exhibits normal growth, L49 appears smaller and forming less leaves which seem to be wilted.

**Figure 3:**
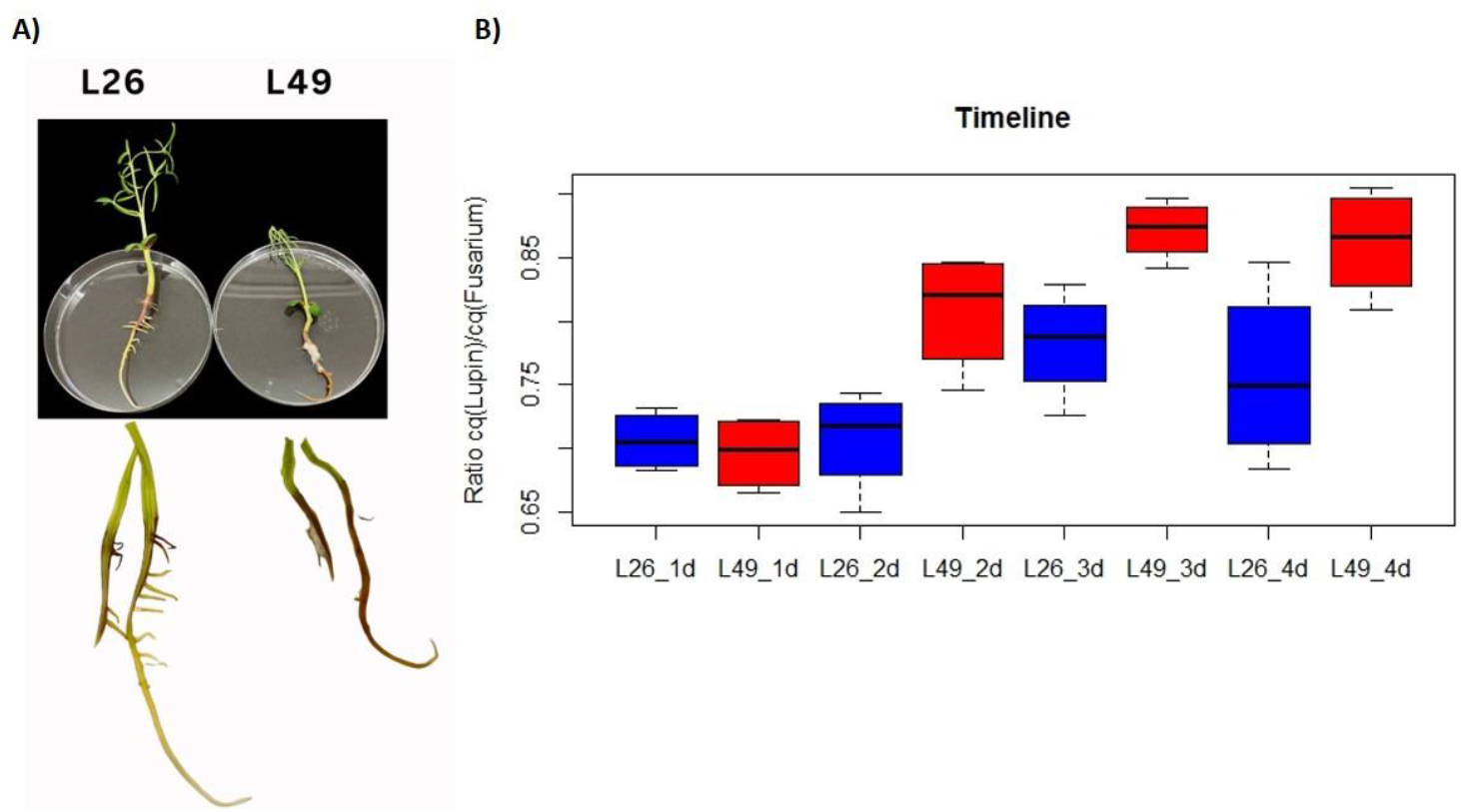
A) L26 and L49 plants and roots 15 dpi; B) calculated cq ratios documenting the infection progress in L26 and L49 over 4 days

### Infection progress over time in the most susceptible accession compared to the most resistant accession

To characterize the infection process in more detail, a time-resolved analysis of the disease progression was carried out. In this context, accessions L26 and L49 were infected and the roots were sampled every day over a total time period of four days. After DNA extraction and qPCR the ratios indicated a slower progress of infection in L26 supporting the previous results (Figure 3B).

While after one day there is no difference between the accessions, there is a significant difference between L26 and L49 (p = 0.019) already two days post inoculation. The differences in the disease progression are further significant after three days (p = 0.011) and after four days (p = 0.045). Taking the phenotypic appearance of the whole plants and the roots into account which were documented 15 days after infection, there is a clear difference between the two accessions. While L49 looks smaller, there is also growth of mycelium on the roots visible. A closer comparison of the roots indicates that the spread of Fol is restricted in L26 while the root of L49 is already completely brownish after 15 days, which is in line with the screening results.

## Discussion

Lupins are emerging, highly valuable crops for sustainable agriculture due to their high protein content and symbiotic nitrogen fixation potential. The development of disease-resistant varieties is essential for minimizing the need of plant protection interventions and at the same time to ensure the economic value for the grower. In this study, we assessed the resistance levels of 20, in this respect yet uncharacterized, NLL accessions. We identified different resistance levels of the accessions towards Fol infection. In previous projects, other characteristics of several of the screened accessions such as yield, lime tolerance, pod number, pod dropping and pod shattering were already evaluated (Eickmeyer 2023). In the accession, L26, that showed the highest Fol resistance in our screen, several desired traits have been identified. L26 is tall, forming large seeds with a high protein content. Even though these traits were confirmed several times by high heredity, a detailed genetic and genomic analysis is still required. An additional trait which was observed at least once was the stability but further data collection is required as well as analysis of heredity. Table 1 lists the accessions tested in our screens for which other known characteristics have been analyzed previously.

**Table 1:** Summary of characteristics of the screened accessions (Eickmeyer 2023, Zeise 2018)

| Accession | Previous characterization |
| --- | --- |
| L26 | High protein content; Tall growing; stable |
| L46 | Lime tolerant, better performance at pH 8 |
| L54 | Good seed quality; eventually reduced susceptibility to Anthracnose |
| L57 | High protein content |
| L58 | eventually reduced susceptibility to Anthracnose |
| L64 | eventually reduced susceptibility to verticillium wilt |
| L66 | High seed quality |
| L80 | High protein content; Tall growing; stable |

The genomes of several *Lupinus angustifolius* accessions have been sequenced and based on these results, a pan-genome has been assembled (Garg et al. 2022). Yet, there are many accessions available which have neither been sequenced nor characterized. Hence, germ plasm banks are a valuable resources offering a broad range of genetic diversity which is an important source for identifying resistance/ susceptibility related genes and further introduce them into crop breeding programs (Mondal, Kumar, and Gnanesh 2023). Sequencing and the evaluation of the genomes of several of our tested accessions is in progress.

Genetic studies have revealed that resistant lupin genotypes possess two dominant, non-allelic resistance genes, *RFO1* and *RFO2*, while susceptible genotypes either lack both or carry only one in a heterozygous state (Jørnsgård et al. 2007). Further research on the resistance genes *RFO1* and *RFO2* against *F. oxysporum* in *Arabidopsis thaliana* has shown that their expression in susceptible ecotypes restricts pathogen growth in the roots. These genes uniquely protect *Arabidopsis* against various *F. oxysporum formae speciales* of the pathogen. *RFO1* encodes a wall-associated kinase-like protein (*WAKL22*), and *RFO2* encodes a receptor-like protein. Transcriptional data indicate only weak gene induction and suppression of many genes, including the Ethylene Response Factor72 (*ERF72*) gene, which plays a role in suppressing programmed cell death and contributes to increased resistance (Chen et al. 2014). It has been suggested that *F. oxysporum* interferes with ethylene signaling since *Arabidopsis thaliana* with a mutated ethylene receptor ETR1 show a higher resistance upon *F. oxysporum* infection (Pantelides et al. 2013). While ethylene in plants is produced via 1-Aminocyclopropane-1-carboxylic acid (ACC) employing an ACC-oxidase, in *F. oxysporum f. sp. tulipae* an alternative pathway where 2-oxoglutarate is converted to ethylene using the ethylene forming enzyme has been identified (Hottiger and Boller 1991). While salicylic acid signaling is contributing to a reduced susceptibility of the plant, jasmonic acid, ethylene, abscisic acid and auxin signaling are potential targets hijacked by *F. oxysporum* (Di, Takken, and Tintor 2016).

It is important to consider that, apart from the genetic background, also environmental factors contribute to the resistance of a plant against pathogens. Previous tests of 547 accessions in four different locations revealed that the resistance is not only depending on the accession but also on environmental factors (Kurlovich, Korneichuk, and Kiselev 1995). Therefore, a complex screening scheme starting with high-content laboratory screens to identify resistance under defined conditions, to genomic analysis of the most contrasting lines up to field experiments under different environmental conditions are required to unequivocally determine if a certain accession may be suitable for further breeding programs.

## Conclusion

In this project, a fast-screening method for NLL resistance towards Fol was developed and the resistance levels of 20 NLL accessions were determined. Among them, we identified one highly susceptible and one highly resistant accession which were confirmed in different additional phenotypic and molecular assays. These two most contrasting lines will be fully sequenced and used for further genomics analysis in an attempt to identify genes that may be crucial for Fol resistance and can then be further characterized on the molecular and biochemical level. The ultimate goal is to breed sweet lupin lines which are resistant to the major pathogens under any environmental conditions and at the same time carrying a large number of additional traits that make them an even more attractive crop plant for sustainable agriculture.

## Author contributions

CR: Method development, screening, data evaluation; EM: method establishment, data evaluation; KT: timeline, data evaluation; TS: experimental design, method establishment, data evaluation, draft manuscript preparation, funding acquisition

